# Alternative splicing detection workflow needs a careful combination of sample prep and bioinformatics analysis

**DOI:** 10.1101/005546

**Authors:** M. Carrara, J. Lum, F. Cordero, M. Beccuti, M. Poidinger, S. Donatelli, R. A. Calogero, F. Zolezzi

## Abstract

**Background:** RNAseq provides remarkable power in the area of biomarkers discovery and disease stratification. The main technical steps affecting the results of RNAseq experiments are Library Sample Preparation (LSP) and Bioinformatics Analysis (BA). At the best of our knowledge, a comparative evaluation of the combined effect of LSP and BA was never considered and it might represent a valuable knowledge to optimize alternative splicing detection, which is a challenging task due to moderate fold change differences to be detected within a complex isoforms background.

**Results:** Different LSPs (TruSeq unstranded/stranded, ScriptSeq, NuGEN) allow the detection of a large common set of isoforms. However, each LSP also detects a smaller set of isoforms, which are characterized both by lower coverage and lower FPKM than that observed for the common ones among LSPs. This characteristic is particularly critical in case of low input RNA NuGEN v2 LSP.

The effect on statistical detection of alternative splicing considering low input LSP (NuGEN v2) with respect to high input LSP (TruSeq) on statistical detection of alternative splicing was studied using a benchmark dataset, in which both synthetic reads and reads generated from high (TruSeq) and low input (NuGEN) LSPs were spiked-in. Statistical detection of alternative splicing (AltDE) was done using prototypes of BA for isoform-reconstruction (Cuffdiff) and exon-level analysis (DEXSeq). Exon-level analysis performs slightly better than isoform-reconstruction approach although at most only 50% of the spiked-in transcripts are detected. Both isoform-reconstruction and exon-level analysis performances improve by rising the number of input reads.

**Conclusion:** Data, derived from NuGEN v2, are not the ideal input for AltDE, specifically when exon-level approach is used. It is notable that ribosomal depletion, with respect to polyA+ selection, reduces the amount of coding mappable reads resulting detrimental in the case of AltDE. Furthermore, we observed that both isoform-reconstruction and exon-level analysis performances are strongly dependent on the number of input reads.

## Background

The application of next-generation sequencing (NGS) to transcriptomics analysis, namely RNAseq, has allowed many advances in the characterization and quantification of transcripts. Recently, several developments in RNAseq methods have provided an advance in the complete characterization of RNA molecules [1]. These developments include improvements in transcription start site mapping, strand-specific measurements, gene fusion detection, small/long RNA characterization and detection of alternative splicing events. [1]. More recently further advances in the application of RNAseq were focused to RNA sequencing approaches allowing RNA quantification from very small amounts of cellular materials or even single cells [2-6]. The main technical steps affecting the outcomes of RNAseq experiments are Library Sample Preparation (LSP) and Bioinformatics Analysis (BA). NGS applications require specific LSP in which fragmented DNA or cDNA molecules are fused with adapters amplified by PCR and sequenced [7]. Since different LSP can have significant impacts on downstream analysis and interpretation of RNAseq data [8], it is evident that robust library preparation methods that produce a representative, non-biased source of nucleic acid material from the genome under investigation are critical. Nevertheless, it has become clear that LSPs contain biases that compromising the quality of NGS datasets can lead to erroneous interpretations [7]. The LSPs now available on the market are various, but they can be organized in two main classes: i) unstranded (high or low input total RNA) and ii) stranded (polyA+ selected or rRNA depleted).

As introduced before, the choice of LSPs does not represent the only critical step in RNAseq. Indeed, the sequence data generated need to be converted into transcript information, e.g. transcript structure, transcript quantification, etc., this step requires an accurate selection of the methodologies for bioinformatics and statistical analysis. The BA for the detection of differentially expressed transcripts are characterized by multiple steps [9] that can have strong influence on the final results. BA can be divided in two categories: i) isoform-reconstruction based differential expression, ii) exon-based differential expression. This work focuses on the definition of the limits and the strengths of LSP as well as the effect of BA on statistical detection of alternative splicing (AltDE). In detail, we investigated the effect of different LSPs (NuGEN v2, TruSeq unstranded/stranded, ScriptSeq) as well as the effect of polyA+ selection versus ribosomal depletion on isoform level detection. Furthermore, we compared NuGEN low input protocol with standard TruSeq protocol in AltDE using prototypic BA for isoform-reconstruction (Cuffdiff) and exon-level analysis (DEXSeq).

## Results

### Library sample preparation (LSP) effects on isoforms detection and isoforms characteristics

We observed how high (100-1500 ng), low (0.5-2 ng) input protocol, polyA+ selection and ribosomal RNA depletion affect isoforms detection. Specifically, we analysed the LSP effect on isoforms coverage/FPKM, exons and exon-exon junctions counts. Total RNA, extracted from the D1 mouse dendridic cell line, was split in aliquots and converted in libraries using the following sample preparation kits: NuGEN v2, ScriptSeq v1, TruSeq unstranded/stranded. The input of total RNA was 0.5 (nu05), 2 (nu2) and 100 ngs (nu100) for NuGEN v2 (nu), 1500 ng for ScriptSeq v1 (ss), 100 (ts100) and 1000 ng (ts1000) for TruSeq unstranded (ts) and 100 ng for TruSeq stranded (tss). All above-mentioned LSPs were performed after polyA+ selection, but the TruSeq stranded LSP, which was also used in association with the ribo-zero ribosomal RNA depletion method (tss_total). For each experimental condition (n05, nu2, nu100, ss, ts100, ts1000, tss, tss_total) 80 million paired-end reads were collected. Reads were mapped against the mouse genome version 9 (mm9). Mapped reads were associated with their corresponding transcript using UCSC annotation and cufflinks [10], as prototypic method for isoform quantification. For each experimental condition we retained only the transcripts characterized by FPKM > 0.1 and average coverage > 0 (Table 1). To define the number of common transcripts detected by the various LSPs we used as reference ts100 detected transcripts. Ts100 was selected as reference because 100 ng total RNA input represents the quantity that can be obtained from a wide range of biological samples, e.g. cell lines, animal model tissues, biopsies, etc..

**Table 1.** Number of isoforms detected using Cufflinks [10] starting from 80 million reads generated by different Library sample preparation.

| Names | KIT | selection | input (ng) | Isoforms (FPKM > 0.1, Coverage > 0) | sample prep specific transcripts with respect to ts100 | common with respect to ts100 | % in common with ts100 |
| --- | --- | --- | --- | --- | --- | --- | --- |
| nu05 | NuGEN | polyA | 0.5 | 21707 | 2597 | 19110 | 77.37 |
| nu2 | NuGEN | polyA | 2 | 24410 | 3502 | 20908 | 84.64 |
| nu100 | NuGEN | polyA | 100 | 25901 | 4012 | 21889 | 88.62 |
| ss | ScriptSeq | polyA | 1500 | 24135 | 2078 | 22057 | 89.30 |
| ts1000 | TruSeq<br>unstranded | polyA | 1000 | 24857 | 1165 | 23692 | 95.92 |
| ts100 | TruSeq<br>unstranded | polyA | 100 | 24701 | 0 | 24701 | 100.00 |
| tss | TruSeq<br>stranded | polyA | 100 | 24318 | 1766 | 22552 | 91.30 |
| tss_total | TruSeq<br>Total | Ribo-Zero | 100 | 11993 | 464 | 11529 | 46.67 |

#### Effect of polyA + selection versus rRNA depletion

All LSPs allow the detection of a similar number of transcripts (Table 1) except for tss_total, generated using total RNAs upon ribosomal depletion, where the number of transcripts is reduced to 46% approximately. In tss_total, the number of mapped reads was in the same order of magnitude of the other experiments. However, the relative amount of coding polyA+ transcripts is diluted, since the input material contains also non-coding RNAs, resulting in a lower sampling of coding transcripts. The above concept is reinforced by the observation that transcripts undetectable in tss_total are those characterized by low coverage/FPKM distributions in ts100 (Fig. 1), as instead the transcripts detected both by tss_total and ts100 show similar coverage and FPKM distributions (Fig. 1).

**Fig. 1.**
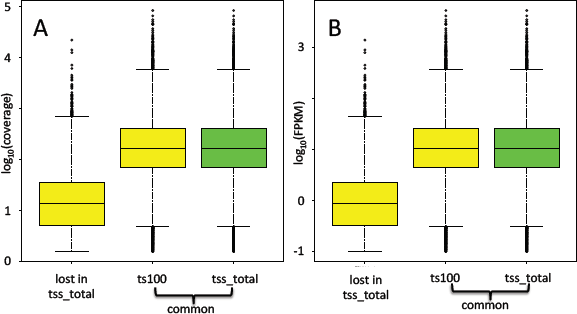
Coverage and FPKM of isoforms in tss_total. A) Coverage distributions. B) FPKM distributions. The majority of the lost isoforms in tss_total are characterized by low coverage and low FPKM.

#### Transcripts detection in low input protocol

We observed that the number of detected transcripts, in NuGEN v2, dependents on the amount of the input material (Table 1). The number of transcripts in common with the ts100 increases moving from 0.5 to 100 ng of total RNA input. Moreover, the number of NuGEN specific transcripts (Table 1, Fig. 2A) increases. The coverage of NuGEN detected transcripts (Fig. 2B, yellow and green boxes) is lower than ts100 detected transcripts (Fig. 2B, violet boxes). This is particularly true for NuGEN specific transcripts (Fig. 2B, yellow boxes). However, the behaviour observed for the coverage does not apply to FPKM distribution (Fig. 2C). Unless for nu05 dataset, NuGEN detected transcripts show FPKM distribution (Fig. 2C, yellow/green boxes) similar to that observed for ts100 dataset (Fig. 2C, violet boxes).

**Fig. 2.**
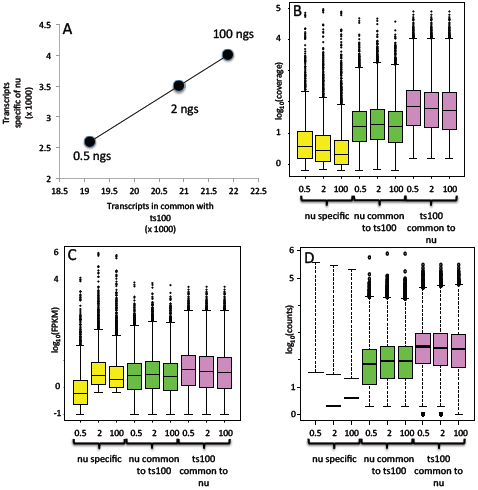
Transcripts coverage and FPKM in low input LSP. A) The number of the LSP specific transcripts increases linearly with the increment of the transcripts in common with ts100 LSP, which depends on total RNA input. B) Coverage for transcripts detected only by low input LSP (yellow boxes) is much lower than the coverage of transcripts in common with ts100 (green boxes). The increment on total RNA input does not improve the coverage for transcripts in common with ts100 (green boxes). Coverage in ts100 LSP has higher coverage than the one obtained by low input LSP. C) Unless for 0.5 ng in low input LSP (yellow), the FPKM of all conditions show a similar distribution. D) The counts of the exons associated with the transcripts in B/C indicate a very low exon counts distribution for the nu05, nu2 and nu100 specific-exons (yellow boxes) and a lower number of exon counts associated to transcripts in common with ts100 in nu05, nu2 and nu100 (green boxes) with respect to ts100 exon counts (violet boxes).

We analysed the coverage and FPKM distributions for ss, tss and ts1000 with respect to ts100 (Fig. 3). The coverage and FPKM distributions of transcripts in common between ss, tss, ts1000 and ts100 are very similar to each other. On the other side the LSP specific transcripts are always characterized by very low coverage/FPKM distributions (Fig. 3). Thus, the low coverage for LSP transcripts in common with ts100 is only a peculiarity of NuGEN derived data.

**Fig. 3.**
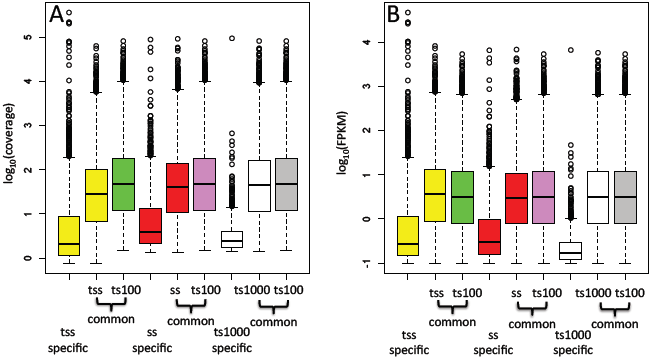
Transcript coverage and FPKM in LSPs with 100-1000 ngs input of total RNA. A, B, C) Coverage for transcripts detected only by tss, ss, ts1000 LSPs is much lower than the coverage of transcripts in common with ts100. Transcripts detected in common with respect to ts100 show nearly identical coverage.

We further investigated this point analysing the raw count distribution for exons belonging both to the transcripts detected by NuGEN and for those transcripts in common with ts100 (Fig. 2D). From this analysis it is clear that exons, belonging to transcripts detected by NuGEN, are characterized by low exon coverage (Fig. 2D, black boxes). This is particularly true for the nu05 sample, where the mean of its exon-counts distribution is not shown since the majority of the exons have 0 counts (Fig. 2D, black boxes). Instead, a mean value less than 10 counts is observed in samples nu2 and nu100 (Fig. 2D, green/violet boxes). In the case of exons detected both by nu and ts100 the exon counts distribution is lower in nu05, nu2, and nu100 (Fig. 2D, green boxes) with respect to ts100 (Fig. 2D violet boxes). The presence of lower coverage for transcripts/exons detected by NuGEN could represent a critical issue in isoform differential expression, since it might affect the bioinformatics quantification of the transcripts/exons.

Finally we also checked the presence of detectable differences in the numbers of exon-exon junction in transcripts specific for nu05, nu2 and nu100 with respect to those in common with ts100 (Fig. 4). The exon-exon junction counts distribution is narrow for transcripts identified using the NuGEN LSP with respect to TruSeq LSP (Fig. 4A). In case we consider the average detection ratio of exon-exon junctions this is lower in NuGEN LSP with respect to TruSeq LSP (Fig. 4B). Considering only isoform-specific exon-exon junctions, i.e. exon-exon junctions allowing discrimination between different isoforms, the differences in average detection ratio become negligible for nu05 and nu2 (Fig. 4B).

**Fig. 4.**
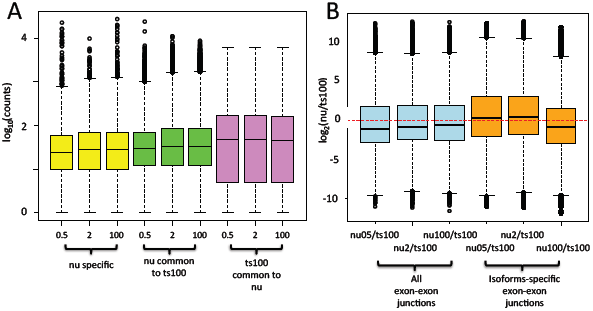
Characteristics of exon-exon junctions in low input LSP. A) The number of counts associated with exon-exon junctions in nu05, nu2 and nu 100 both for LSP-specific transcripts and for those transcripts in common with ts100 have a detection range, which is narrow with respect to those detectable with ts100. B) Log_2_ ratio between nu05, nu2, nu100 and ts100 exon-exon junction counts, for transcripts detected in common by the two LSPs (light blue boxes); log_2_ ratio between nu05, nu2, nu100 and ts100 transcripts-specific exon-exon junction counts (orange boxes).

### Benchmark datasets

The observations provided in the previous paragraph highlight that NuGEN v2 has different characteristics with respect to high input LSPs (TruSeq unstranded/stranded, ScriptSeq). NuGEN protocol using 0.5 ng of input total RNA (nu05) has a very limited ability (-23% with respect to ts100, Table 1) to detect isoforms with respect to TruSeq unstranded protocol using 100 ng input total RNA (ts100). The isoform detection with NuGEN protocol using 2 (nu2) or 100 (nu100) ng still remains a little less efficient of ts100 with respect to the other LSPs. Although nu100 looses, with respect to ts100, only 12% of the detected transcripts (Table 1) it will not be used in standard experiments because of the higher complexity/cost of the protocol compared to other LSPs requiring the same input quantity of RNA. Nu2 represents the best compromise between the need of a low input RNA quantity and the number of detected isoforms (-16% with respect to ts100, Table 1). Therefore, we decided to compare the effect of nu2 and ts100 on the detection of differential isoform expression by BA approaches. To address this question we created benchmark datasets where nu2 and ts100 reads were spiked-in, within a common background made of TruSeq unstranded data reads (C1-C5 ts1000, T1-T5 ts100; Fig. 5, Additional Table 1S). Specifically, we spiked-in reads derived from 20, 40 and 80 million reads of both nu2 (NU20/40/80 datasets, Additional Table 2S) and ts100 dataset (TS20/40/80, Additional Table 2S). With this design, we obtained an isoform-level differential expression between C and T groups for 27 transcripts (Fig. 6). Furthermore, to investigate the dependency of the BA approaches on gene-specific isoforms complexity we constructed a synthetic dataset where complex expression composition of isoforms for the same gene was also present (Additional Table 3S). Synthetic reads were characterized by having a uniform distribution over all transcripts and 58 differentially expressed transcripts between C and T groups were generated (Fig. 5, 6).

**Fig. 5.**
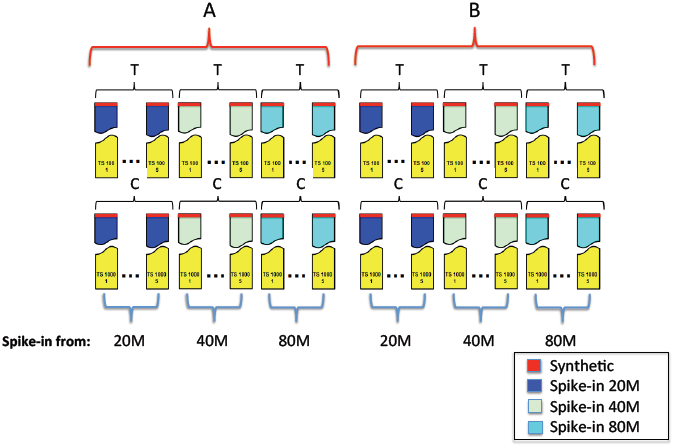
Benchmark dataset. A) Three datasets (TS20, TS40, TS80), based on spike-in of TruSeq (input: 100 ng total RNA) reads extracted respectively from 20, 40 and 80 million reads were generated using a common background made by 5 different TruSeq library preps having as input 100 ng total RNA (T) and 5 different TruSeq library preps having as input 1000 ngs total RNA (C). B) Three datasets (NU20, NU40, NU80), based on spike-in of NuGEN (input: 2 ngs total RNA) reads extracted respectively from 20, 40 and 80 million reads were generated using the common background described above. Synthetic spike-ins are present both in A and B.

**Fig. 6.**
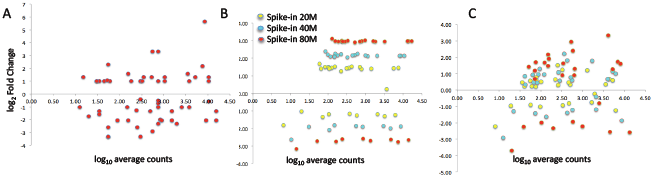
Differential expression of the spike-in data. A) Synthetic spike-ins characterized by uniform coverage over the transcripts. B) TruSeq spike-ins. C) NuGEN spike-ins.

### Isoforms differential expression analysis

The identification of differentially expressed isoforms was investigated on the above mentioned datasets using the following approaches: cuffdiff [11], as prototypic for isoforms-reconstruction approaches, and DEXSeq [12], as prototypic for exon-level analysis. The increase of the number of reads also increases the detection of differentially expressed isoforms independently by the dataset under analysis, i.e. NU or TS (Fig. 7A). In the case cuffdiff we used two available versions of the program, cuffdiff 1 and cuffdiff 2. Cuffdiff 2, with respect to cuffdiff 1, also embeds the estimation of the over-dispersion due to biological replications [11]. Cuffdiff 1 detects a fixed number of transcripts independently by the number of the reads used to generate the spike-in on the TS dataset (Fig. 7A, blue bar). Otherwise, on the NU dataset the differential expression detection increases on the basis of the number of reads used in the spike-in generation. It is notable that using 80 million reads cuffdiff 1 detects the same number of alternative spliced transcripts discovered using 20 million reads in the TS dataset. Thus, alternative splicing events detection efficacy of cuffdiff 1 seems to be quite inefficient if NU datasets are used.

**Fig. 7.**
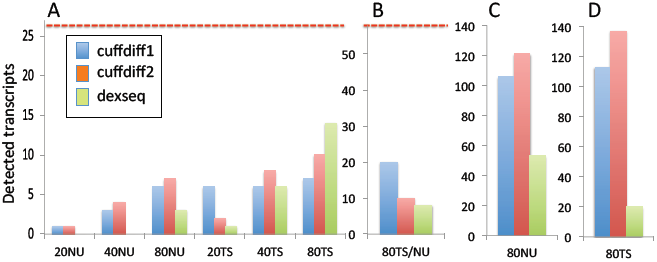
Statistical detection of spliced isoforms. A) True positive transcripts detected as differentially expressed between C1-C5 and T1-T5 groups as function of the spike-ins (20,40,80) and of the LSP (NU, TS). B) True positive synthetic transcripts detected as differentially expressed between C1-C5 and T1-T5. Only the 80 millions spike-in is considered since the synthetic spike-ins are identical over all the datasets. C) False positive transcripts detected in the dataset NU80, depending on the BA used in the analysis. Only the 80 millions spike-in is considered since the false positive are nearly in the same amount for 20, 40 and 80 millions spike-ins. D) False positive transcripts detected in the dataset TS80 depending on the BA used in the analysis. Only the 80 millions spike-in is considered since the false positive are nearly in the same amount for 20, 40 and 80 millions spike-ins.

In case of cuffdiff 2, there is an increment in the detection of differentially expressed transcripts correlated to the number of reads used to generate the spike-in; this is observable in both TS and NU datasets (Fig. 7A, orange bar). Cuffdiff 2 detects a greater number of differentially expressed transcripts than cuffdiff 1 in both the datasets except in the case of 20 million reads TS (Fig. 7A, blue and orange bars).

The exon-level analysis by DEXSeq is quite inefficient with respect to both versions of cuffdiff in the case of the NU dataset (Fig. 7A, green bar) and in general in the samples characterized by a low number of input reads. However, in case of TS80, where spike-ins are derived by 80 million reads, DEXSeq detects the highest number of alternative spliced isoforms (Fig. 7A green bar). The overall detection of differentially expressed spike-in data using DEXSeq on TS80 reaches approximately the 50% of the total true positive isoforms. It is notable that the overall false positive detection rate is particularly low in the case of DEXSeq (Fig. 7C).

We also evaluated the level of overlaps between results obtained with isoforms-reconstruction approaches and exon-level analysis (Fig. 8). In the case of NU80 dataset the overlap is minimal, probably because of the poor performances of DEXSeq on NuGEN spike-in data (Fig. 8A). On the other side DEXSeq has in common with both versions at least 66% of the detected true positive transcripts in the TS dataset (Fig. 8B).

**Fig. 8.**
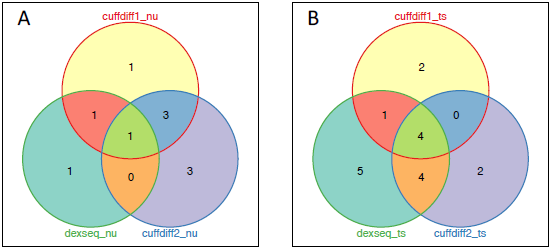
Overlap of results among different BA in isoforms detection. A) NU80 data set. B) TS80 dataset.

The experiments performed on the synthetic dataset reveal inferior detection efficiency (Fig. 7B). The best results are obtained by cuffdiff 1 detecting approximately 34% of the total true positive isoforms.

## Discussion

In this paper we present the first comparative evaluation of the combined effect of Library Sample Preparation and Bioinformatics Analysis on alternative splicing detection.

Library Sample Preparations both stranded and non-stranded preparations (ss, tss, ts), working with at least 100 ng of total input RNA and undergoing polyA+ enrichment, show a similar behaviour for commonly detected transcripts. Considering the comparison between polyA+ selection versus ribosomal depletion the reads sampling is distributed between coding and non coding transcripts, hence, the transcripts detection is significantly impaired for the low expressed transcripts.

Transcripts that are specifically detected only by a LSP show poor coverage and they are probably very little informative for isoform detection, because of the non-uniform count coverage at exon-level. In the case of NuGEN, low input protocol, the number of LSP-specific transcripts increases with the rise of the amount of total RNA input. However, those LSP-specific transcripts are characterized by low coverage and in general by very low exon-level counts. FPKM estimation for those transcripts can be misleading since it has a behaviour very similar to that observed for the transcripts in common with TruSeq protocol. Nu05, nu2 and nu100, even for the transcripts in common with ts100, show a lower coverage and exon counts distribution with respect to those obtained with TruSeq LSP (ts100). The experiments on benchmark datasets reveal that the lower exon counts generated from NU datasets (NU20/40/80) negatively affect the ability of exon-level based approach (DEXSeq) to detect alternative splicing events. On the other side, in case high number input reads are used and the preparation is done using TruSeq protocol, i.e TS80 dataset, exon-level based approach provides the best results. Its notable that the overlap in alternative spliced isoforms is only partial between isoforms-reconstruction approaches, and exon-level analysis. Exon-level analysis detects a higher number of true positive alternative splicing transcripts with a lower number of false positive with respect to isoforms-reconstruction approach.

## Conclusions

Our results indicate that a low input protocol, as NuGEN v2, is not suitable for alternative splicing analysis due to the limited coverage at exon-level. Furthermore, the performances of both isoforms-reconstruction approaches, and exon-level analysis are in general comparable. However, it is notable that in case of high number of input reads the exon-level analysis provides a higher detection rate of alternative splicing events with a reduced level of noise.

## Materials and Methods

### RNA isolation and purification

Total RNA was extracted from D1 mouse cell line [13]. Total RNA was extracted with Trizol Reagent (Invitrogen) followed by RNeasy micro clean-up procedure (Qiagen) as per manufacturer’s instructions. Total RNA integrity was assessed by Agilent 2100 Bioanalyzer (Agilent) and the RNA Integrity Number (RIN) was calculated; RNA sample had a RIN = 9.5.

### Library Preparation

#### Illumina TruSeq RNA

D1 total RNA was mixed with ERCC RNA Spike in Control Mix 1 (Ambion) and subjected to library preparation using Illumina TruSeq RNA Sample Preparation kit version 1 (Low Sample Protocol) with slight modifications. Briefly, polyA containing RNA molecules were purified from the D1 RNA samples using polyT oligo-attached magnetic beads. Thermal fragmentation followed after two rounds of enrichment for polyA+ mRNA. cDNA was synthesized from the RNA fragments using reverse transcriptase (Superscript II) and random primers. This was followed by second strand cDNA synthesis and the resulting dsDNA went through an end repair process, the addition of a single A base, and ligation of the adapters. The products were then purified and enriched with 12 cycles of PCR to create the cDNA library. Two additional rounds of purification of the cDNA libraries were done using Agencourt Ampure XP SPRI beads (Beckman Courter) to remove >600 bp double stranded cDNA. The length distribution of cDNA libraries was monitored using DNA 1000 kits on the Agilent Bioanalyzer. All libraries were subjected to an indexed PE sequencing run of 2 × 51 cycles on an Illumina HiSeq 2000.

#### NuGEN Ovation RNA-Seq system version 2 – Dedicated Read Barcode

Total RNA was processed for cDNA synthesis using Ovation RNA-Seq system version 2 (NuGEN Technologies) according to the manufacturer’s protocol. ERCC RNA spike in control Mix 1 were mixed with D1 RNA before cDNA synthesis. Briefly, first strand cDNA synthesis was performed using a unique first strand DNA/RNA chimeric primer mix and reverse transcriptase (RT). A DNA/RNA heteroduplex double-stranded cDNA was generated by fragmentation of the mRNA within the cDNA/mRNA complex allowing the DNA polymerase to synthesize a second strand. The DNA then underwent SPIA amplification. SPIA is an isothermal linear amplification in which RNase H is used to degrade RNA in the DNA/RNA heteroduplex at the 5’ end of the first cDNA strand, after which the SPIA primer binds and DNA polymerase then initiates replication at the 3’ end of the primer, displacing the existing forward strand. The process of SPIA DNA/RNA primer binding, DNA replication, strand displacement and RNA cleavage is repeated, resulting in rapid accumulation of SPIA cDNA. SPIA cDNA from each replicate were sheared to get a size range of 25 bp to 400 bp with the bulk of the material at 150 bp. This was done by sonication (Covaris model S2) with duty cycle 10, intensity 5 and cycle/burst 100 for 300 s. 200 ng of the sheared DNA were then used for library preparation using the Encore NGS Multiplex System 1 (NuGEN Technologies) according to manufacturer’s protocol where the fragmented DNA underwent end repair to generate blunt ends, adaptor ligation (with 4 bases indexing tags) and amplification to enrich the fragments with ligated adapter sequences. The resultant libraries were then purified using the Agencourt RNAClean XP beads. Additional round of purification were done with Agencourt Ampure XP SPRI beads (Beckman Courter) and the libraries were eluted in 20 ul. 4 ul of each purified library underwent 10 cycles of PCR amplification using Illumina TruSeq PCR reagents. All libraries were subjected to an indexed PE sequencing run of 2 × 51 cycles on an Illumina HiSeq 2000.

#### Epicentre ScriptSeq v1

PolyA containing mRNA molecules were purified from 1.5 ug D1 total RNA using polyT oligo-attached magnetic beads. cDNA libraries were prepared from the resultant polyA+ RNA and ERCC RNA spike in control Mix 1 were added.

The RNA samples were chemically fragmented using the StarScript Reverse Transcriptase Buffer and the cDNA Synthesis Primer was annealed to the RNA. 5′ end-tagged cDNA (equivalent to the 3′ end of the original RNA) was produced by random-primed cDNA synthesis. This was followed by 3′-Terminal Tagging of the cDNA using the Terminal-Tagging Oligo (TTO) which randomly annealed to the cDNA, including to the 3′ end of the cDNA and served as template for the extension of the cDNA by DNA polymerase. The resulting di-tagged cDNAs (at both their 5′ and 3′ ends) were tagged purified using Qiagen MinElute PCR Purification Kit (Qiagen). Enrichment of the purified di-tagged cDNAs were done with 12 cycles of PCR and libraries were purified with Agencourt AMPure XP SPRI beads (Beckman Courter) according to the standard protocol. Two additional rounds of Agencourt Ampure XP beads were conducted to remove > 600 bp double stranded cDNA. ScripSeq dedicated read barcode design, which is analogous to TruSeq barcoded adapters, were used for the samples. All libraries were subjected to an indexed PE sequencing run of 2 × 51 cycles on an Illumina HiSeq 2000.

#### Illumina TruSeq Stranded Total RNA

D1 total RNA were mixed with ERCC RNA Spike in Control Mix 1 and subjected to library preparation using Illumina TruSeq Stranded Total RNA Sample Preparation kit (Low Sample Protocol) with slight modification. The removal of ribosomal RNA was done using Ribo-Zero Gold rRNA removal beads which deplete samples of both cytoplasmic and mitochondrial ribosomal RNA. After depletion, the RNA was purified and fragmented into small pieces using divalent cations using thermal fragmentation. First strand cDNA synthesis was performed using reverse transcriptase (Superscript II) and random primers from the cleaved RNA fragments. This was followed by second strand cDNA synthesis using DNA Polymerase I and RNase H where second strand was generated with dUTP in place of dTTP. These blunt end cDNA fragments had the addition of a single A base and subsequent ligation of the adapter. The products were purified and enriched with 12 cycles of PCR to create the final cDNA library. Two additional rounds of purification of the cDNA libraries were done using Agencourt Ampure XP SPRI beads (Beckman Courter) to remove > 600bp double stranded cDNA. All libraries were subjected to an indexed PE sequencing run of 2 × 51 cycles on an Illumina HiSeq 2000.

#### Illumina TruSeq Stranded mRNA

D1 total RNA was mixed with ERCC RNA Spike in Control Mix 1 and subjected to library preparation using Illumina TruSeq Stranded mRNA Sample Preparation kit (Low Sample Protocol) with slight modification. Briefly, polyA containing mRNA molecules were purified from the D1 RNA samples using polyT oligo-attached magnetic beads. Thermal fragmentation followed after two rounds of enrichment for polyA+ mRNA. First strand cDNA synthesis was performed using reverse transcriptase and random primers from the cleaved RNA fragments. This was followed by second strand cDNA synthesis using DNA Polymerase I and RNase H where the second strand was generated with dUTP in place of dTTP. These blunt end cDNA fragments then had the addition of a single A base and subsequent ligation of the adapter. The products were purified and enriched with 12 cycles of PCR to create the final cDNA library. Two additional rounds of purification of the cDNA libraries were done using Agencourt Ampure XP SPRI beads (Beckman Courter) to remove > 600 bp double stranded cDNA. All libraries were subjected to an indexed PE sequencing run of 2 × 51 cycles on an Illumina HiSeq 2000.

### Spike-in dataset

The common background of the spike-in dataset was made using paired-end reads generated preparing with the TruSeq unstranded protocol 5 libraries, starting with 1000 ng of total RNA extracted from the D1 cell (C1-C5), and 5 libraries starting with 100 ng of total RNA D1 cells (T1-T5) (Additional Table 1S). The true positive set (TP) of transcripts was defined in the following way: exon counts for samples C1-C5 and T1-T5 were loaded in R using DEXseq package [12] and UCSC mm9 annotation (28232 genes). Genes characterized by at least three isoforms were selected (6582). Then, those genes having at least one transcript characterized by at least one exon discriminating it from the other isoforms were selected (6313). The genes were further filtered, removing all transcripts characterized by having, for the discriminating exons, less than 10 counts in total in C1-C5 and T1-T5 samples (2970). Out of the 2970 transcripts 27 were randomly selected and from them one of the isoform was used for spike-in experiment (Additional Table 2S). Sequences data, generated with NuGEN v2, using as input 2 ng input total RNA, and with TruSeq unstranded, using 100 ng input total RNA, were used to construct three datasets made respectively of 20, 40 and 80 million reads. Each dataset was mapped against the mm9 mouse genome and the reads mapping to the 27 transcripts were extracted and spike-in C1-C5, T1-T5 to simulate transcripts up and down-regulation within two experimental conditions (Additional Table 2S).

### Synthetic dataset

Out of the 2970 transcripts described in the previous paragraph we randomly selected 58 transcripts. For each transcript we decide to spike-in a specific number of reads (Additional Table 3S). Then, 10000 values from a normal distribution, having as mean the defined number or reads to be spiked-in (Additional Table 3S) and a standard deviation of 1/10^th^ of the mean, were generated. A value was then randomly selected and used to define the number spike-in to be placed in C1-C5 and T1-T5. Then, within each transcript, a uniform distribution of sequence start points made of 10000 elements was defined. From this uniform distribution we selected the number of reads to be spiked-in. Synthetic pair-end reads 2 × 51 nts were constructed. Reads were associated with a quality score of 40 and used to generate fastq files, which were added to C1-C5 and T1-T5 background samples.

### Isoforms quantification and statistical detection of alternative spliced isoforms

Nu05, nu2, nu100, ts100, ts1000, ss, tss, tss_total, C-1-C5 and T1-T5 fastq data were mapped with STAR [14]. For Nu05, nu2, nu100, ts100, ts1000, ss, tss and tss_total isoform quantification was done with cufflinks [15]. Exon-level quantification was done using DEXSeq [12] and exon-exon junction quantification was done with subjunc function of the Rsubread [16] Bioconductor package. Cuffdiff 1 and cuffdiff 2 [11], prototypic BA based on isoform-reconstruction, were used for detection of alternative spliced isoforms between C-1-C5 and T1-T5 groups using mm9 UCSC annotation. Isoforms were considered differentially expressed if characterized by q-value ≤ 0.05. For exon-level analysis was used DEXSeq [12]. Isoforms were considered differentially expressed if at least one isoform-specific exon was detected as differentially expressed between C-1-C5 and T1-T5 groups with a Benjamini & Hochberg adjusted p-value ≤ 0.05.

## Authors’ contributions

CM generated spike-in datasets, LJ made all LSPs, PM performed NGS data QC and mapping, CF executed the bioinformatics comparisons between different LSPs, BM executed the bioinformatics comparisons between different LSP, DS supervised the bioinformatics analysis, CRA investigated the effects of LSPs on BA and wrote the paper, ZF supervised the sample preparations and wrote the paper.

## Acknowledgements

We wish to thank Prof. Paola Ricciardi-Castagnoli for providing D1 cells. This study was funded by grants from the Epigenomics Flagship Project EPIGEN, MIUR-CNR, European Seventh framework program, Health.2012.1.2-1, NGS-PTL grant n. 306242 and by the core fund of Singapore Immunology Network, Agency for Science, Technology and Research (A*STAR), Singapore.

## Additional files

**Additional Table 1S.** Background Pair-end reads datasets

|  | **C1** | **C2** | **C3** | **C4** | **C5** | **T1** | **T2** | **T3** | **T4** | **T5** |
| --- | --- | --- | --- | --- | --- | --- | --- | --- | --- | --- |
| **Background PE reads** | 13.4M | 25.8M | 10.9M | 13.4M | 10.0M | 8.4M | 7.4M | 19.6M | 16.1M | 7.3M |

**Additional Table 2S.** Endogenous and spike-in counts

|  | **C1-C5 mean** | **nu2** |  |  | **ts100** |  |  | **T1-T5 mean** | **nu2** |  |  | **ts100** |  |  |
| --- | --- | --- | --- | --- | --- | --- | --- | --- | --- | --- | --- | --- | --- | --- |
| **isoform** | **endogenous** | **20M** | **40M** | **80M** | **20M** | **40M** | **80M** | **endogenous** | **20M** | **40M** | **80M** | **20M** | **40M** | **80M** |
| **uc009txr.2** | 755 |  |  |  |  |  |  | 607 | 159 | 318 | 541 | 973 | 2047 | 4074 |
| **uc009ghp.2** | 1378.4 |  |  |  |  |  |  | 1068.4 | 1419 | 2637 | 9710 | 1768 | 3641 | 7228 |
| **uc009gho.2** | 2038 | 1368 | 2600 | 10043 | 2623 | 5301 | 10618 | 1566 |  |  |  |  |  |  |
| **uc007jtu.1** | 5453.2 |  |  |  |  |  |  | 4618 | 2781 | 5523 | 10820 | 816 | 16015 | 31665 |
| **uc007jtv.2** | 6860.4 |  |  |  |  |  |  | 5656.6 | 2940 | 5830 | 11437 | 10013 | 19594 | 38859 |
| **uc007gin.1** | 2863.8 | 489 | 974 | 2082 | 4208 | 8428 | 16800 | 2476 |  |  |  |  |  |  |
| **uc008mib.2** | 5186 |  |  |  |  |  |  | 4281.6 | 1263 | 2539 | 6094 | 7315 | 14629 | 29243 |
| **uc009eup.1** | 5863 | 7945 | 15397 | 29280 | 7934 | 15681 | 31240 | 4609.6 |  |  |  |  |  |  |
| **uc007wpv.1** | 659.6 |  |  |  |  |  |  | 521.8 | 202 | 367 | 756 | 951 | 1814 | 3585 |
| **uc007wps.1** | 546.8 | 398 | 722 | 2001 | 752 | 1492 | 3024 | 449.8 |  |  |  |  |  |  |
| **uc012baj.1** | 385 | 346 | 628 | 1088 | 482 | 1000 | 1915 | 280.6 |  |  |  |  |  |  |
| **uc008egx.1** | 358.6 |  |  |  |  |  |  | 261.4 | 362 | 657 | 1135 | 457 | 944 | 1795 |
| **uc012hbj.1** | 299.6 |  |  |  |  |  |  | 234.2 | 135 | 261 | 494 | 396 | 814 | 1578 |
| **uc007dpl.2** | 104.4 | 106 | 219 | 416 | 134 | 261 | 530 | 78 |  |  |  |  |  |  |
| uc009oda.1 | 117.2 |  |  |  |  |  |  | 93.8 | 15 | 33 | 259 | 166 | 329 | 638 |
| uc007mki.1 | 142.4 |  |  |  |  |  |  | 109.8 | 49 | 105 | 198 | 206 | 413 | 767 |
| uc007mkf.1 | 92.6 |  |  |  |  |  |  | 77.4 | 14 | 30 | 47 | 142 | 292 | 540 |
| uc009fyx.1 | 4.4 | 16 | 29 | 52 | 11 | 23 | 35 | 4 |  |  |  |  |  |  |
| uc007aui.1 | 100.4 |  |  |  |  |  |  | 75.6 | 55 | 109 | 267 | 134 | 256 | 515 |
| uc012ffj.1 | 107.6 |  |  |  |  |  |  | 81.4 | 41 | 76 | 180 | 135 | 261 | 546 |
| uc009gyv.2 | 65.4 | 56 | 108 | 192 | 101 | 212 | 370 | 50.6 |  |  |  |  |  |  |
| uc007hwe.1 | 70.2 |  |  |  |  |  |  | 64.6 | 31 | 58 | 83 | 118 | 214 | 436 |
| uc009rfh.2 | 244.4 |  |  |  |  |  |  | 178 | 339 | 653 | 1187 | 255 | 558 | 1150 |
| uc009rfi.1 | 20.4 | 19 | 29 | 77 | 21 | 55 | 115 | 17.6 |  |  |  |  |  |  |
| uc009nfj.1 | 399.2 |  |  |  |  |  |  | 306.2 | 75 | 150 | 267 | 582 | 1155 | 2178 |
| uc009mpg.1 | 61.2 |  |  |  |  |  |  | 51 | 9 | 14 | 117 | 85 | 185 | 350 |
| uc007ris.1 | 54.2 |  |  |  |  |  |  | 37.8 | 15 | 31 | 63 | 85 | 163 | 289 |

**Additional Table 3S.**
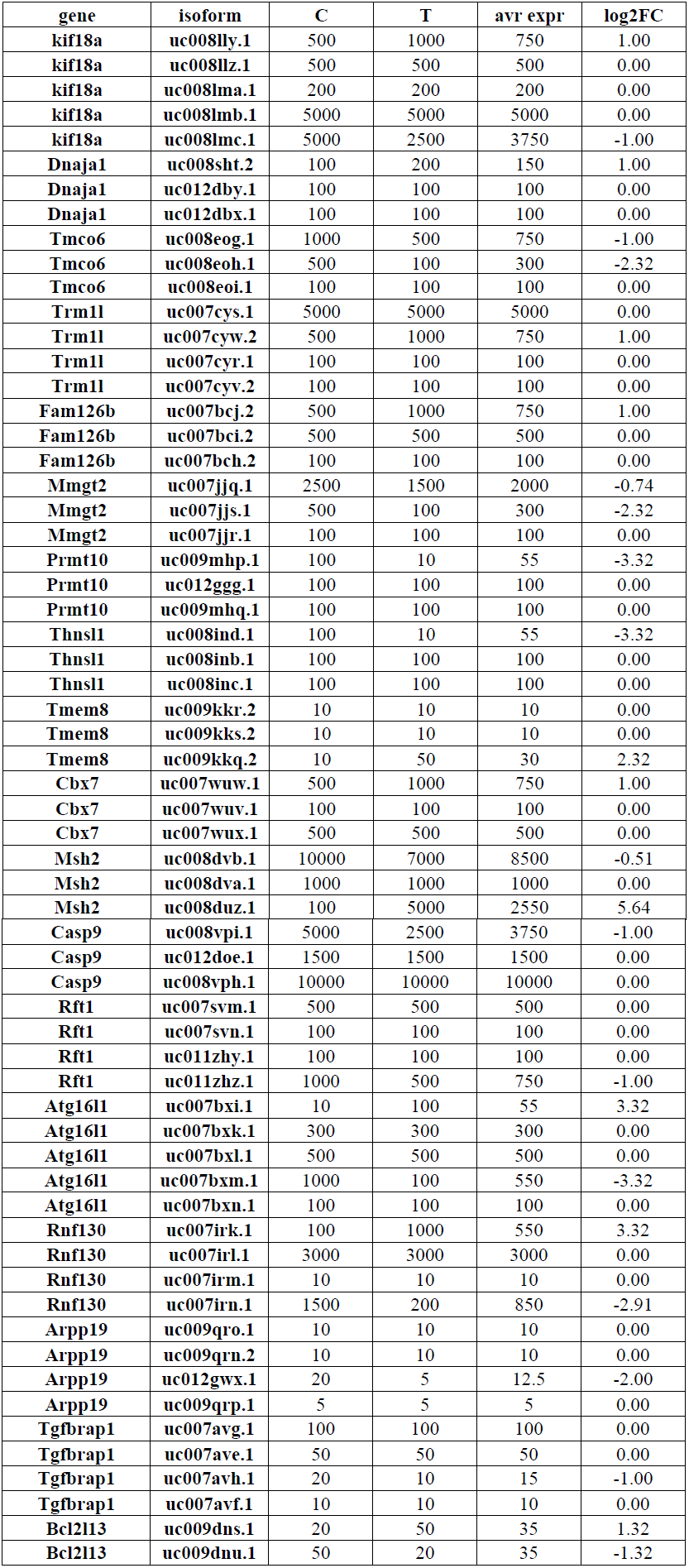
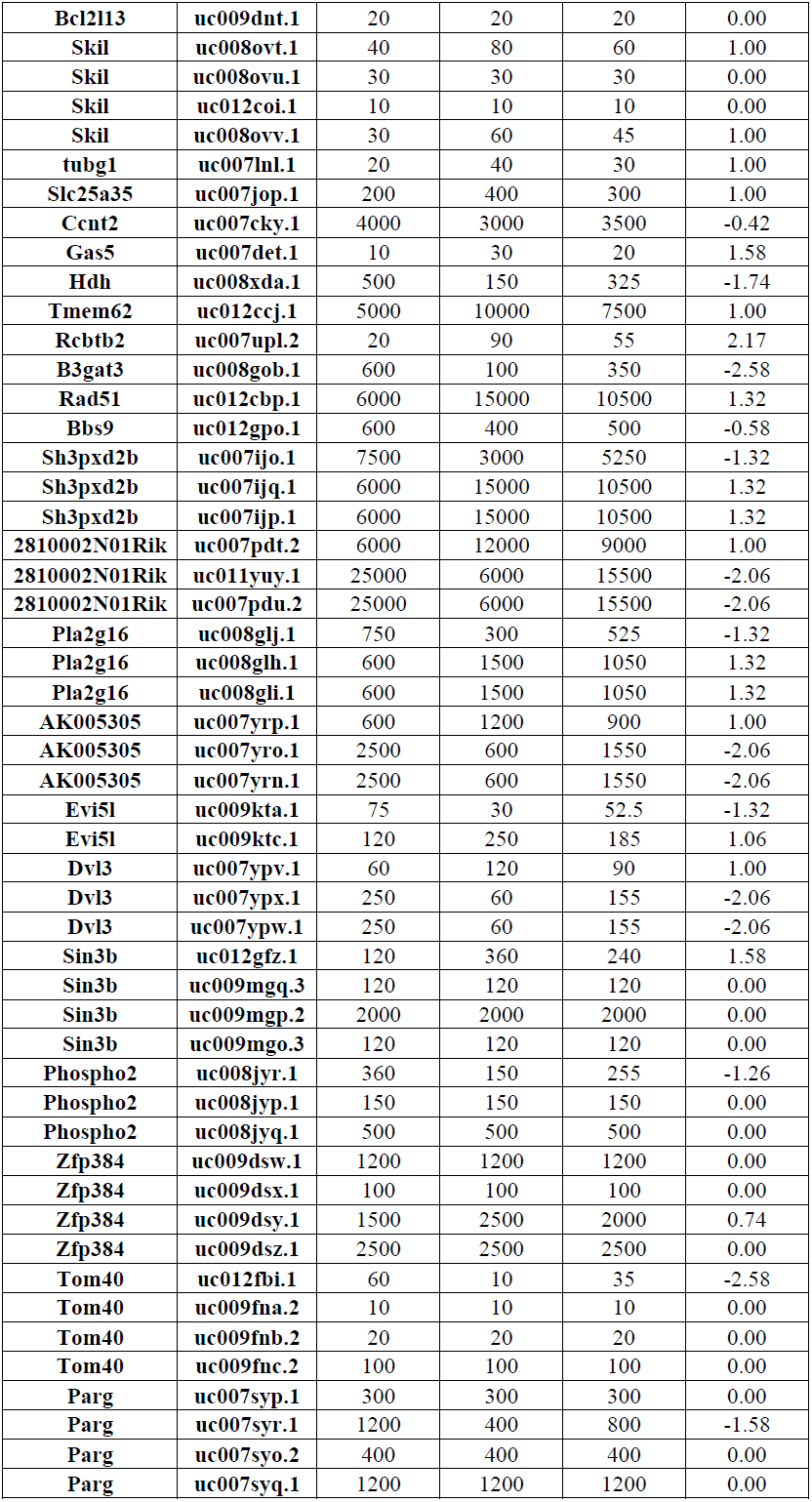
Synthetic spike-in data

